# Attraction to pheromones in *Caenorhabditis elegans* can be reversed through associative learning

**DOI:** 10.1101/476648

**Authors:** M. Dal Bello, A. Pérez-Escudero, F. C. Schroeder, J. Gore

## Abstract

Despite the ubiquity and importance of chemical signaling, we have only limited insight about the role of learning in the response to pheromones. Here, we demonstrate that responses to pheromones can be reprogrammed through associative learning. In particular, we show that attraction to ascaroside pheromones in the model nematode *Caenorhabditis elegans* can be reversed by training the animals to associate either a pheromone blend or single synthetic ascarosides with the lack of food. This behavioral plasticity alters worm preference for pheromones following consumption of a food patch, possibly improving foraging in natural environments. By bridging the gap between the current knowledge on the chemical language and the learning abilities of *C. elegans*, we provide insight on the possible links between learning and chemical signaling in animals.

---

Chemical signals underlie the behavior of all organisms, from bacteria to humans. Although examples of chemical communication were known since the ancient Greeks, it is in 1959, after the discovery of the silk moth sex cue bombykol, that Karlson and Lüscher (*1*) introduced the term “pheromones”. Pheromones are molecules that are externally secreted by an organism and act by triggering a specific reaction, such as a stereotyped behavior and/or a developmental process, in another individual of the same species (*1*). Among other things, pheromones regulate mate selection, promote aggregation, and aid in locating food or avoiding predators; in many species, they constitute the basis for social interactions. Responses induced by pheromones are typically predictable and innate meaning that they do not require specific learning (*2*).

The nematode *Caenorhabditis elegans* has an extensive chemical language that relies on more than 150 different molecules belonging to the family of ascarosides (*3*). Since several signal transduction pathways are conserved among invertebrates, the study of pheromone signaling in *C. elegans* has greatly contributed to the understanding of intraspecific chemical communication. By signaling population density, ascaroside pheromones regulate worm development, entry into dauer (a developmentally arrested larval stage) (*4–8*) and the plasticity in the response to both attractive and repulsive compounds (*9, 10*). Ascarosides are also potent attractants for adult worms, some of them mainly drawing males, others hermaphrodites (*11–16*).

Attraction to ascarosides, in all instances, has been assumed to be innate (*11*–*14*, *16*, *17*). Nevertheless, *C. elegans* exhibits a high level of behavioral plasticity (*18*), being able to learn and remember environmental stimuli (smells, tastes, oxygen levels and temperature) that anticipate attractive or aversive signals (*19–21*). In particular, worms can learn to associate these stimuli with the presence of bacterial food, and the result of this learning exercise is that worms are attracted through chemotaxis to that particular stimulus (*20*).

Here we demonstrate that response to ascarosides is another case of associative learning. Attraction can indeed be turned into repulsion by training *C. elegans* to associate either a pheromone blend produced by worms or single synthetic ascarosides to the absence of food. We also explore the possible ecological consequences of the observed behavioral plasticity in the response to pheromones, highlighting that changing preference for pheromones following bacterial food consumption may help the worms to forage more efficiently in their natural environment.

## Results

We first asked what is the response of a *C. elegans* natural isolate (MY1) to the pheromone cocktail it produces, as this simulates what worms are exposed to in their natural environment (*22*). We found that a natural pheromone blend obtained by collecting and filtering the supernatant of a worm liquid culture (*23, 24*) is a potent attractant. In a standard chemotaxis assay (*20, 25*), the number of worms arriving to a spot containing the pheromone blend was more than twice the number arriving at a spot containing a control solvent (Fig. 1A). By contrast, the chemotaxis-defective mutant PR691 (*26*), which cannot detect chemicals, could not distinguish between the pheromone blend and a control solvent (Fig. 1A). The explanation that has been given to this observation in the past decade is that attraction of the worms to the pheromone blend is an innate response (*11*–*14*, *16*, *17*).

**Figure 1.**
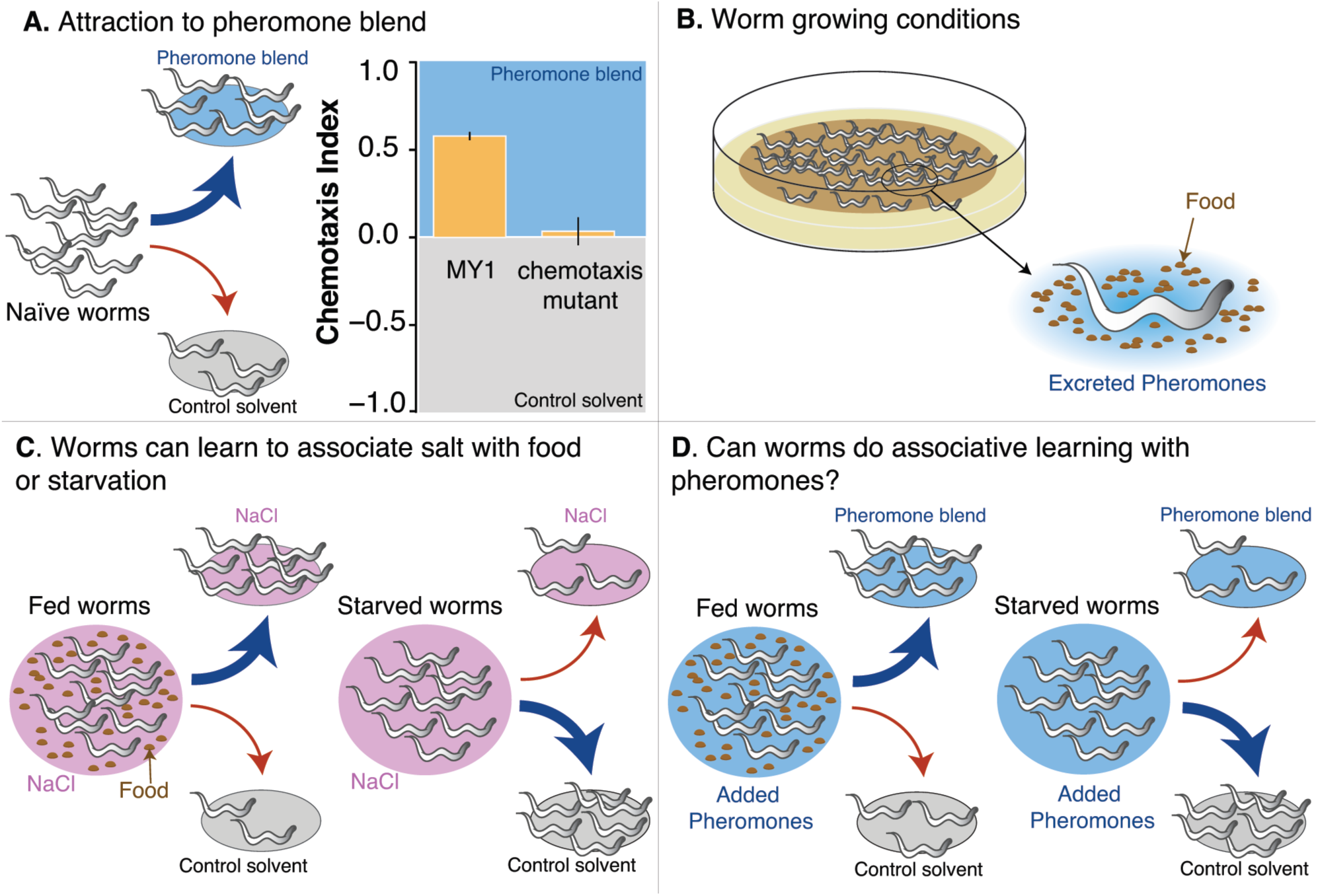
Is *C. elegans* attraction to pheromones a case of associative learning? **A.** Natural isolate MY1 is attracted to a pheromone blend obtained from the supernatant of a MY1 liquid culture. A chemotaxis mutant (PR691), instead, can’t distinguish between the pheromone blend and a control solvent. Attraction is measured with a chemotaxis assay, in which age-synchronized naïve worms are given the choice between the pheromone blend and a control solvent. Chemotaxis index (CI) is defined as (N_P_ –N_C_) / (N_P_+N_C_), where N_P_ and N_C_ are the number of worms at the pheromone and control spot, respectively (mean CI ± SEM, for MY1, n. experiments=4; for PR691, n. experiments =1). **B.** The chemotaxis assay is performed on worms that grow at high density with plenty of food (*E. coli* OP50), excreting and sensing pheromones. **C.** *C. elegans* can learn to associate salt with positive or negative stimuli. Worms fed in the presence of salt (NaCl) are attracted by it, while worms starved in the presence of salt avoid it (*20*). **D.** If associative learning regulates *C. elegans* response to pheromones, attraction to the pheromone blend is due to the positive association with food that the worms make during growth, and it can be reversed by starving the worms in the presence of the pheromone blend.

However, while growing, worms are simultaneously exposed to both bacteria and ascarosides, which are continuously secreted by the animals (*27*) (Fig. 1B). Hence, an alternative explanation to “innate attractive behavior” can be that worms are attracted to the pheromone blend because of the positive association with food that they learn to make during growth on the plate. If associative learning regulates *C. elegans* response to pheromones, this positive association could be reversed just by training the worms to associate the pheromone blend with starvation, as has been demonstrated for salt (Fig. 1C) (*20*). We would therefore expect that animals fed in the presence of pheromones are attracted to them, while worms starved in the presence of pheromones avoid them (Fig. 1D).

We tested this hypothesis in a learning experiment in which MY1 worms were trained for six hours to associate their pheromone blend with either a positive stimulus, bacterial food, or a negative one, starvation (Fig. 2A). We found that chemotaxis to the pheromone blend, which is strong when worms are trained with food and pheromones, is lost by starving worms in the presence of pheromones (Fig. 2B, blue and yellow bars). Worms have learned to associate the pheromone blend with the absence of food and in response they have changed their behavior. This suggests that, in the search for food, worms may avoid regions with pheromones if they experienced starvation in the presence of pheromones.

**Figure 2.**
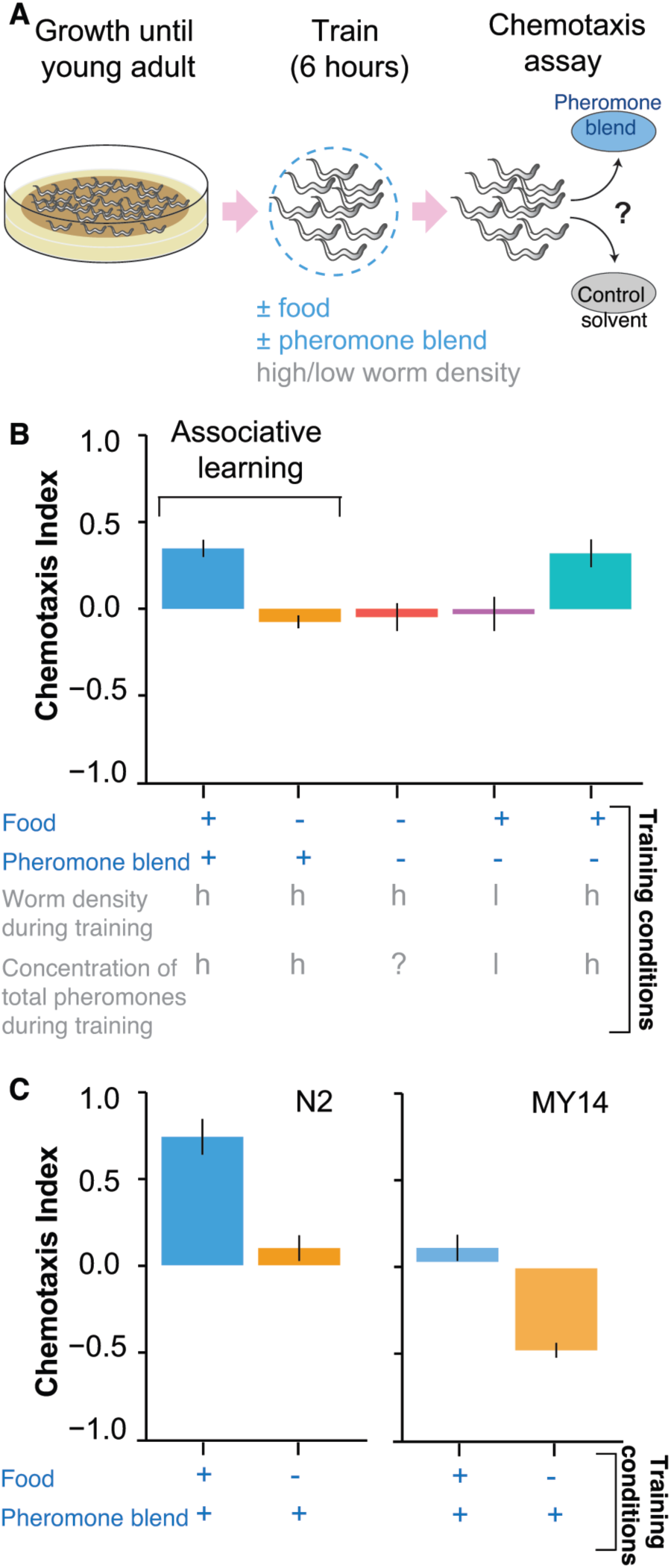
Associative learning regulates *C. elegans* response to pheromone blend. **A**. Worms grow at high density and with plenty of food until young adult. Animals are then transferred to conditioning plates, where they spend 6 hours. Worms are then assayed for chemotaxis to the pheromone blend. **B**. The natural isolate MY1 is capable of associative learning with the pheromone blend. Blue labels along the x axis indicate the training conditions; grey labels indicate instead the density of worms (h, high or l, low) and the total pheromone concentration (added pheromone blend + pheromones excreted by worms during the training period, h, high or l, low) inside the conditioning plates. Chemotaxis index for each different training condition (worms at high density): ‘+food+pheromone blend’, blue bar; ‘–food+pheromone blend’, yellow bar; ‘–food– pheromone blend’, red bar; ‘+food–pheromone blend’, turquoise bar. Purple bar: ‘+food– pheromone blend’ with worms at low density. Mean CI ± SEM, n. experiments= 4. **C**. Also the strain N2 and MY14 are capable of associative learning with the pheromone blend. Chemotaxis index for ‘+food+pheromone blend’ (blue bar), ‘–food+pheromone blend’ (yellow bar), worms at high density (mean CI ± SEM, for N2, n. experiments=1; for MY14, n. experiments=2).

To determine the worm response in the absence of information about pheromones, animals were also trained without pheromone blend, both with and without bacterial food (Fig. 2A). Preference for the pheromone blend is lost when worms are trained in a plate without food nor pheromone blend (Fig. 2B, red bar). We interpret this as meaning that six hours without any information on pheromones is enough to make worms “forget” the positive association between food and pheromones they acquired during growth.

In contrast, preference for the pheromone blend is not lost when worms are trained in a plate in which we added food but not the pheromone blend (Fig. 2B, turquois bar). In this condition, animals, whose density in conditioning plates is high, are simultaneously exposed to the food, which we provided, and the pheromones, which they are excreting (*10*). Hence, they may still be learning to associate pheromones with food. If this is true, worms trained with just food would lose chemotaxis towards the pheromone blend only in a condition in which their density is low enough to prevent the pheromone concentration from building up during the conditioning period. Consistent with this hypothesis, the preference for the pheromone blend goes to zero when animals are trained with food at low density (Fig. 2B, purple bar). The fact that worms lose the positive association between food and pheromones they learned during growth provides further support that *C. elegans* attraction to pheromones is not just innate.

A possible concern is that our observation of associative learning can be explained if starved worms simply behave differently than fed worms, losing the preference for the pheromone blend. However, our experiments training the worms in the presence of food but without added pheromones at high or low worm density show that it is not only the feeding state of the worm that determines the response to the pheromone blend, since equally fed worms showed attraction at high density and no preference at low density. This result further reinforces our hypothesis that learning can modify worm response to pheromones.

We next asked whether associative learning with pheromones is a general phenomenon in *C. elegans*. To address this question, worms of the lab strain N2 and of another natural isolate, MY14, were trained for six hours to associate the pheromone blend with either bacterial food or starvation. We found that associative learning occurs in both strains, regardless of the magnitude of the attraction of each strain to the pheromone blend (Fig. S1). In particular, chemotaxis to the pheromone blend is strongly reduced in N2 worms, which displays strong attraction, and reversed in MY14 worms, which instead shows only a slight attraction to the pheromone blend (Fig. 2C). Differences in the relative strengths of attraction and repulsion to pheromones between the laboratory strain N2 and most wild isolates have been shown to result from a mutation in the neuropeptide-Y receptor locus, *npr-1*(*13*). These results argue that associative learning of pheromone signals is likely common among *C. elegans* strains.

The pheromone blend contains, in addition to a cocktail of ascarosides, other products of worm metabolism, compounds deriving from the decomposition of dead worms and bacteria, and perhaps other unknown substances. To test whether altered responses to ascaroside pheromones underlie the observed learning, we repeated the experiment with pure synthetic ascarosides. We tested two synthetic molecules – ascr#5 and the indole ascaroside icas#9 – at the concentration at which each pheromone was attractive for MY1 strain (Fig. S2). Ascr#5 is abundantly produced by *C. elegans* (*15*) and can act synergistically with other ascarosides promoting dauer formation (*7*). Unlike other dauer-inducing ascarosides, ascr#5is attractive for hermaphrodite N2 individuals (*13*). Icas#9 exerts attraction on hermaphrodites, promoting aggregation (*14*), and suppresses exploratory behavior by slowing down worms while they are crawling on food (*16*).

To test for associative learning with pure ascr#5 and icas#9, worms were trained to associate each ascaroside to either bacterial food or starvation and then assayed for chemotaxis to the ascaroside they were trained with (Fig. 3). Worms fed in the presence of either ascr#5 or icas#9 prefer the ascaroside while this preference is reversed when worms are starved in the presence of either of the two ascarosides (Fig. 3). Analogously to the experiment with the pheromone blend, six hours without exposure to ascarosides are enough to lose the positive association between food and pheromones (Fig. 3). These results support our hypothesis that the worms were learning from the pheromones in the crude pheromone blend. We concluded that through associative learning, *C. elegans* is able to reverse its response to pheromones.

**Figure 3.**
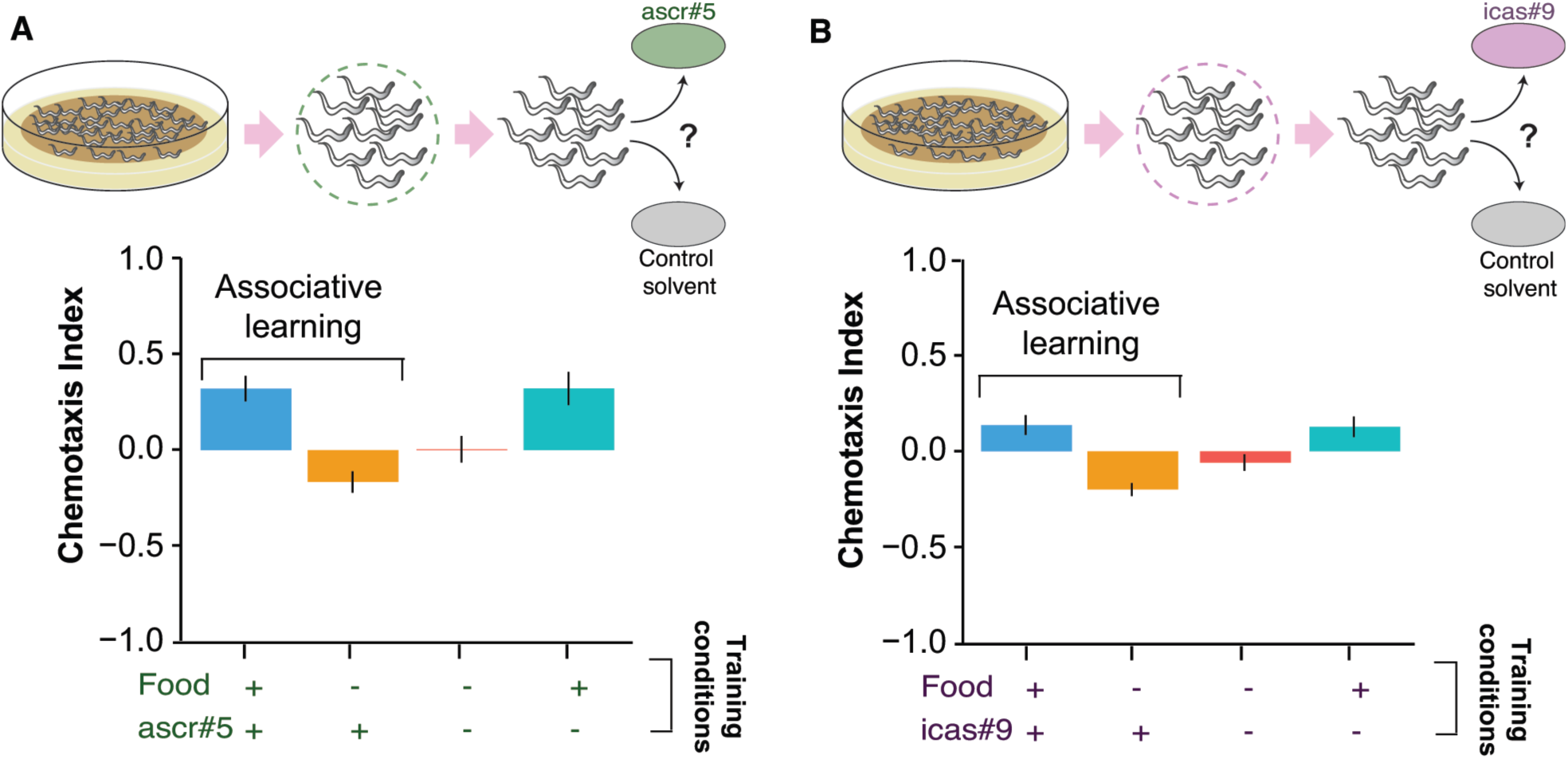
Associative learning can reverse the MY1 response to pure ascarosides (panel A ascr#5, panel B icas#9). Worms grow at high density and with plenty of food until young adult. Animals are then transferred to conditioning plates, where they spend 6 hours training with either ascr#5 or icas#9. Worms are then assayed for chemotaxis to the ascaroside they were trained with. Chemotaxis index (mean CI ± SEM, for ascr#5: n. experiments =3; for icas#9: n. experiments =5) for the different training conditions (indicated in green and violet along the x axis, all at high worm density): ‘+ food + ascr#5 / icas#9’ (blue bars, positive association: food and ascarosides), ‘–food + ascr#5 / icas#9’ (yellow bars, negative association: starvation and ascarosides), ‘– food – ascr#5 / icas#9’ (red bars), ‘+ food – ascr#5 / icas#9’ (turquoise bars).

A potential confounding effect in the learning experiments could stem from the fact that worms during the training period are still producing and excreting pheromones, possibly interfering with the chemotaxis assay. To rule out this effect, we repeated the learning experiment with ascr#5 using a *C. elegans* strain that doesn’t produce pheromones but does sense them (*14*) (*daf-22* DR476). Similar to our results with the MY1 natural isolate, the *daf-22* worms go to ascr#5 when trained to associate it with food, avoid ascr#5 when trained to associate it with starvation, and lose their preference when trained in the absence of both ascr#5 and food (Fig. S3). These findings further reinforce the fact that the modification in the worm behavior is due to pheromones. Interestingly, *daf-22* mutants go to ascr#5 also when trained with just food (Fig. S3). Intuitively, this could contrast with the idea of worm response to pheromones being completely dependent on learning processes. This result suggests that part of the response of wild type *C. elegans* to pheromones is innate, but also that it can be modified by experience.

Associative learning underlies behavioral plasticity, i.e. the ability to tune behavior depending on the context. Several species exhibit behavioral plasticity as an adaptive response to different environments and as a vital strategy for dealing with environmental variation (*28*). *C. elegans’* optimal habitats, i.e. rotting plant stems, flower and fruits, are ephemeral and randomly distributed in space and time (*29, 30*). To survive, *C. elegans* has evolved a boom-and-bust life cycle, in which worms rapidly proliferate on bacteria-rich rotting material, and as soon as food is depleted they enter a non-feeding stage (dauer larva), which acts as a dispersal mechanism (*30*). However, bacterial food is heterogeneously distributed also at smaller spatial scales, i.e. on the rotting material (*30*) and worms need to make decisions about how to explore it (*31, 32*), integrating information on food availability and pheromones (*22*). Pheromones concentrate around food patches that have been discovered, signaling either presence of food, if worms have just started to consume the food patch, or absence of food, if the food patch has been exploited. If worms were always attracted to pheromones then they would frequently end up in already spent patches. One possible mechanism to avoid this is to mark spent patches, and it is known that worms starved for long time periods secrete a pheromone profile that is different from fed worms (*15, 27*). Another mechanism that could act at shorter time scales is to change response to pheromones. We therefore hypothesize that behavioral plasticity depending on associative learning with pheromones may help worms to forage more efficiently.

We consider a possible scenario of worms exploiting food patches on a rotting fruit (Fig. 4A). Initially, worms leaving a freshly depleted food patch are still likely to find a non-depleted food patch by following pheromone cues. However, after some time most patches will be depleted and worms will benefit from avoiding pheromone cues, since they are likely associated to exhausted food patches. This ultimately would result in a wider search for food (Fig. 4A). As an experimental test, we ran a modified chemotaxis assay in which worms encountered a food patch before choosing between the pheromone blend and the control solvent (Fig. 4B). Under our hypothesis, worm preference for either the pheromone blend or the control solvent should change over time, from the pheromone blend initially to the control solvent later. Consistent with this expectation, worms went to the pheromone blend for the first 3 hours of the experiment and then changed their preference, i.e. they avoided the pheromone blend, by the 5^th^ hour (Fig. 4B). Although further investigation is needed to ascertain the impact of such processes in the wild, these results suggests that plasticity in the behavioral response to pheromones may be a further adaptation to exploit ephemeral food sources in a most efficient manner. Intriguingly, *C. elegans* individuals engage in a primitive form of farming by redistributing bacteria while foraging (*31*), and having navigational cues that help avoid areas where nutrients, necessary for bacterial growth, have already been depleted may increase farming success.

**Figure 4.**
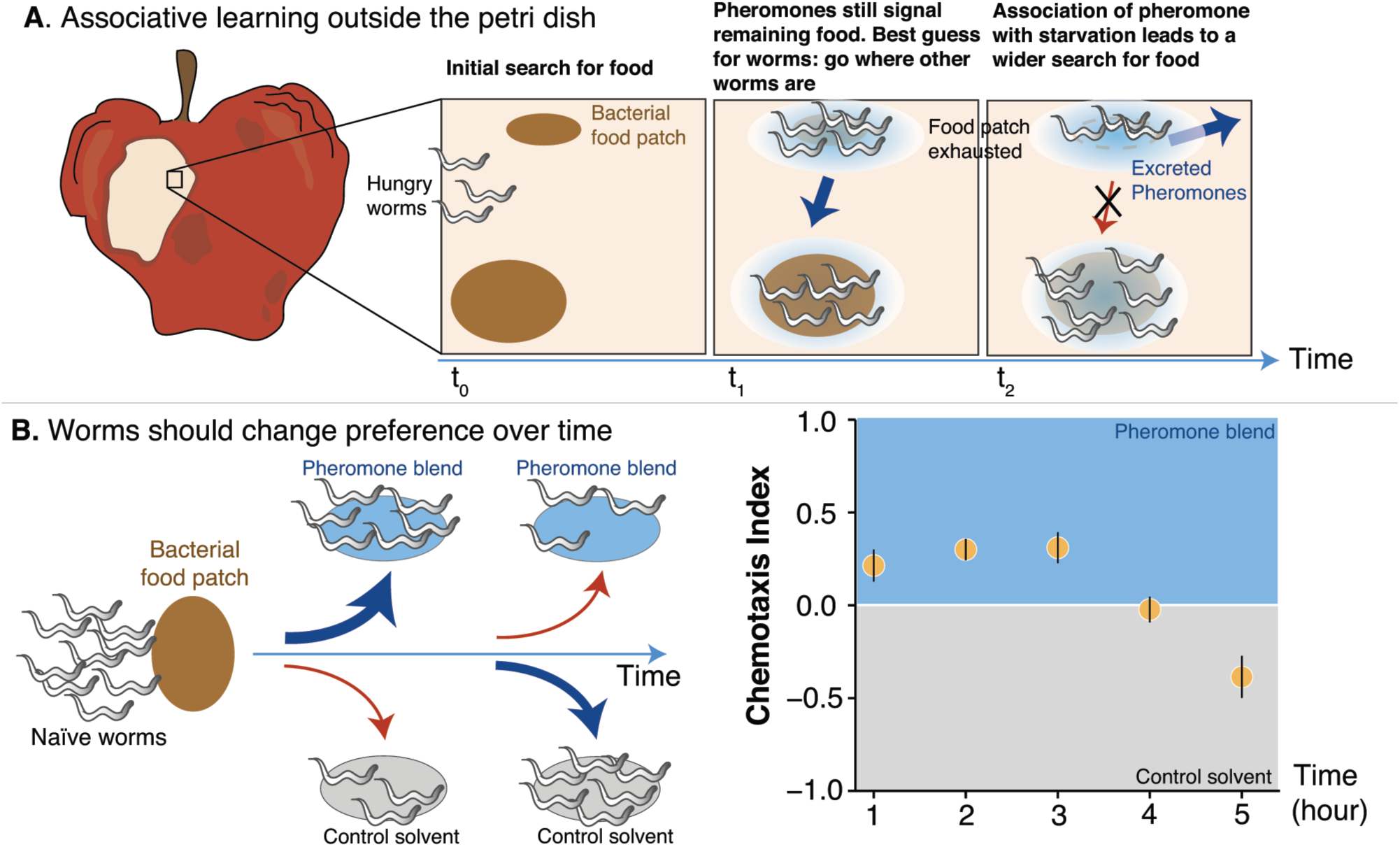
Associative learning with pheromones may help worms to forage more efficiently. **A.** We consider a possible scenario of worms exploiting food patches on a rotting piece of fruit. During the rotting process the bacterial patches could be exhausted at different times. Therefore, worms leaving a freshly depleted food patch are still likely to find a non-depleted food patch by following pheromone cues. However, when most patches become depleted, worms will benefit from avoiding pheromone cues, which are likely to be associated to exhausted food patches. Changing preference for pheromones in time would allow the worms to increase the chances to find unexploited food patches. **B**. Under this hypothesis, the preference for either the pheromone blend or the control solvent of naïve worms presented with a food patch should change over time, from the pheromone blend initially to the control solvent later. In the plot, chemotaxis index calculated on the MY1 worms that, each hour, reach the two spots (mean CI ± SEM, n. experiments =1, n. plates= 25, n. adult worms per plate ~ 80).

## Discussion

Since the discovery of the *dauer* pheromone about 20 years ago, chemical signaling in *C. elegans* has been under ever increasing scrutiny. However, no studies have examined the interplay between learning and chemical signaling in order to elucidate mechanisms underpinning worm behavior. Here we show that attraction to ascarosides is a learned response that depend on the capacity of *C. elegans* to associate environmental stimuli, in this case food, with pheromones.

Exploration of the link between pheromones and behavioral plasticity has recently given a new twist to the research on chemical signaling. In mice and bees, it has been shown that pheromones can influence behaviors that are not the primary target of their action. For example, several compounds present in the urine of male mice become attractive to females only after simultaneous exposure the pheromone darcin (*33, 34*). Similarly, exposure of young worker honey bees to pheromones can modulate learning of aversive associations and reward responsiveness (*35, 36*). As such, pheromones have been proposed as regulators of behavioral plasticity, being the conditioning agent driving the response to chemical compounds with difference valence.

Moreover, chemical cues that regulate important behavioral responses but rely on learning have been discovered (*37–40*). To distinguish them from pheromones they are now identified as signature odors (*38*). For example, the chemical profile of ants displays species-specific molecules, but each colony produces different combinations and ratios of these molecules. Individuals of one colony learn to associate the particular mixture of odors with their nest-mates, helping to avoid conflicts with them while recognizing intruders from a different colony (*37*). Despite the existence of signature odors, this study, to our knowledge, is the first asking whether the response to molecules classified as pheromones can be modified by experience. Whether this applies to other animals than *C. elegans* has not been examined in detail, but these results should prompt more studies investigating the extent to which some innate responses to pheromones can be reprogrammed through experience.

*C. elegans* is one of the best studied animals on earth, which enabled major advances in chemical signaling, behavior, and neuroscience. Our work, demonstrating that *C. elegans* is able to learn to associate pheromones with positive and negative stimuli, suggests that this highly tractable model system could be used to uncover the basis of learning-dependent phenotypic plasticity towards pheromones. Future studies will be required to determine what types of stimuli worms can learn to associate with pheromones, as well as the breadth of behavioral and other phenotypic responses that can be reprogrammed.

## Materials and Methods

### Strains and culture conditions

We used two *Caenorhabditis elegans* strains recently isolated from the wild, MY1 (Lingen, Germany) and MY14 (Mecklenbeck, Germany), as well as N2 (Bristol, UK), the chemotaxis-defective mutant PR691 and the *daf-22* strain DR476. All the strains have been obtained from the Caenorhabditis Genetic Centre (CGC). Worms were grown at 21-23 °C (room temperature) on nematode growth media (NGM) plates (100 mm) seeded with 200 μl of a saturated culture of *E. coli* OP50 bacteria (*41*). As for OP50 culture, a single colony was inoculated into 5ml of LB medium and grown for 24 h at 37 °C.

### Pheromones

We obtained the crude pheromone blend by growing worms in liquid culture for 9 days (at room temperature and shaking at 250 rpm) (*15*). Individuals from one plate were washed and added to a 1-liter flask with 150ml of S-medium inoculated with concentrated *E. coli* OP50 pellet made from 1 liter of an overnight culture. Concentrated *E. coli* OP50 pellet was added any time the food supply was low, i.e. when the solution was no longer visibly cloudy (*41*). The pheromone blend was then obtained by collecting the supernatant and filter-sterilizing it twice. A new pheromone blend was produced every 3 months. We used the pheromone blend produced by MY1 also for tests with N2 and MY14 for two main reasons: to minimize variation among experiments and because the type and amount of ascarosides produced by these strains should be the same (*16*). Pure synthetic ascarosides (ascr#5 and icas#9) were obtained from the Schroeder lab and kept at −20°C in ethanol. Each time an experiment was performed, an aqueous solution at the desired molar concentration was prepared (10 μM for ascr#5 and 10 pM for icas#9). The control solvent for the pheromone blend is S-medium, while the control solvent for the pure ascarosides is an aqueous solution with the same amount of ethanol present in the ascaroside aqueous solution (*14*).

### Chemotaxis assay

Chemotaxis assay has been performed in 60 mm NGM plates, in which worms are given the choice between pheromone (either 20 μl of the pheromone blend or 20 μl of a pure ascaroside in aqueous solution) and a control solvent (20 μl) (*20, 25*). The two spots are deployed ~3 cm apart from each other (Fig. S4A). Shortly before the start of the assay, 1 μl of 0.5 M sodium azide is added to both spots in order to anaesthetize the animals once they reach the spots. Animals, either naïve or trained, are placed equidistant from the two spots and left to wander on the assay plate for 1 hour at room temperature (Fig. S4A). The average number of worms in each experiment is indicated in the figure captions. The assay plates were then cooled at 4°C and the number of worms around each spot was counted using a lens. The chemotaxis index is then calculated as 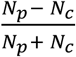, where *N*_*p*_ is the number of worms within 1 cm of the center of the pheromone spot, while *N*_*c*_ is the number of worms within 1 cm of the center of the control spot. The number of independent experiments is indicated in the figure captions. For each experiment, we usually performed 10 replicated assays. The average number of worms used in each replicated assay across all experiments is ~ 50.

### Learning experiment

Worms are grown until they become young adults in NGM plates seeded with 200 μl of saturated *E. coli* OP50 bacteria. Then worms are washed off the plates with wash buffer (M9 + 0.1% triton), transferred to an Eppendorf tube and washed twice by spinning down the worms and replacing the supernatant with fresh wash buffer each time. After that, animals are transferred to conditioning plates where they are trained to associate pheromones (pheromone blend / pure ascaroside) either with a positive stimulus (food) or a negative one (starvation). To induce the positive association, worms trained on an NGM plate seeded with 200 μl of a saturated culture of *E. coli* OP50 blended with 200 μl of pheromone blend or aqueous solution of pure ascaroside (+ food + pheromone). To induce the negative association, worms are conditioned on an NGM plate seeded with 200 μl of pheromone blend or aqueous solution of pure ascaroside (– food + pheromone). As controls, worms are also placed in conditioning plates with 3) nothing (– food – pheromone) and 4) only food (+ food – pheromone). Conditioning plates are prepared ~16 hours before the training starts, so that bacteria can grow and form a lawn. In the case of pure ascarosides, the concentration of ascaroside that was added for the conditioning plate was higher than the concentration at which the worms were tested chemotaxis to compensate for the diffusion of the ascaroside throughout the agar in the conditioning plates. Ascr#5 was added at a concentration of 100 μM onto conditioning plates, while icas#9 was added at a concentration of 1 μM. Worms spent 6 hours on the conditioning plates at room temperature, after which they are assayed for pheromone chemotaxis.

### Modified chemotaxis assay (choice after food)

It is a chemotaxis assay performed on naïve worms that encounter a food patch before making the choice between the pheromone blend and the control solvent. We used 100 mm NGM plates in which we deployed 20 μl of the pheromone blend, 20 μl of control solvent and 15 μl of a diluted OP50 *E. coli* culture at equal distance from each other (Fig. S4B). In the pheromone and control spots, 2 μl of 0.5 M sodium azide was added in order to anaesthetize the animals once they reached the spots. Since the anesthetic action of sodium azide lasts for about 2 hours in this set-up, another 1μl was added two hours after the beginning of the assay in both spots. Naïve animals were placed close to the bacteria spot, so that they stop and consume that patch before chemotaxis towards the two cues. Worms are left to wander freely on the assay plate until the food is finished (5 hours). The number of worms around the two spots was counted every hour and the chemotaxis index was calculated based on the number of new worms that reached the two spots during each hour.

## Supporting information

## Acknowledgments

The authors would like to thank the members of the Gore lab for feedback on the manuscript and Ying K. Zhang for assistance with the synthesis of ascarosides. This work was supported by NIH and the Schmidt Foundation. APE is funded by EMBO Postdoctoral Fellowship Grant ALTF 818- 2014 and Human Frontier Science Foundation Postdoctoral Fellowship Grant LT000537/2015. All *C. elegans* strains were provided by the CGC, which is funded by NIH Office of Research Infrastructure Programs (P40 OD010440).

The authors declare no competing interests.

