## Supplementary material for "Attraction to pheromones in *Caenorhabditis elegans* can be reversed through associative learning"

**This PDF file includes:**

Figs. S1 to S4

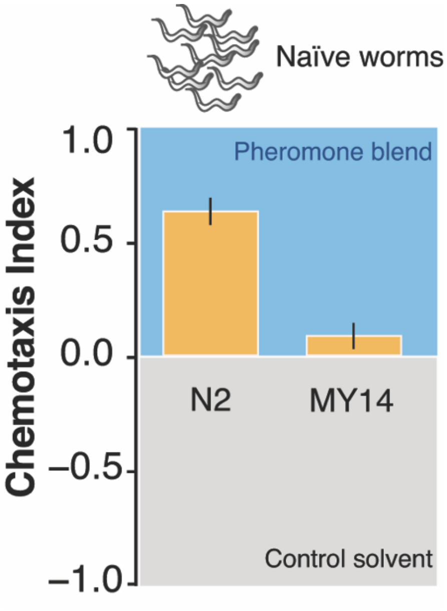


**Fig. S1.** N2 is strongly attracted to a pheromone blend obtained by collecting and filtering the supernatant of a liquid culture of MY1 worms, while natural isolate MY14 is slightly attracted to it. Attraction is measured with a chemotaxis assay performed on age-synchronized naïve worms. Mean CI ± SEM, n. experiments for both N2 and MY14 =2).

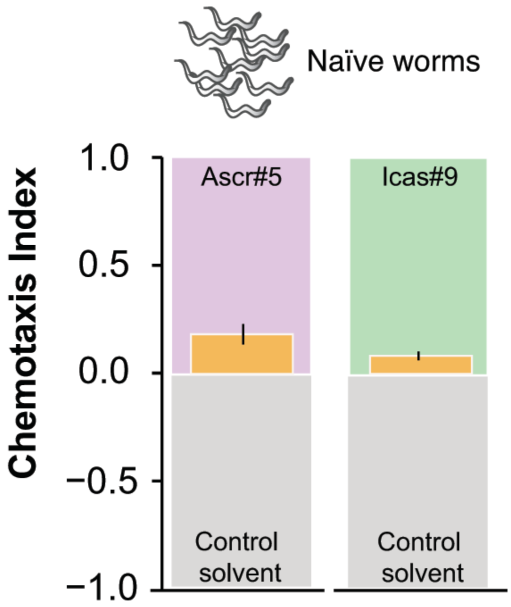


**Fig. S2.** Naïve MY1 worms exhibit modest attraction to synthetic ascarosides. Ascr#5 is attractive at a 10 μM concentration (mean CI ± s.e.m. across plates, n. experiments =2) while icas#9 is attractive at a 10 pM concentration (mean CI ± s.e.m. across plates, n. experiments =3).

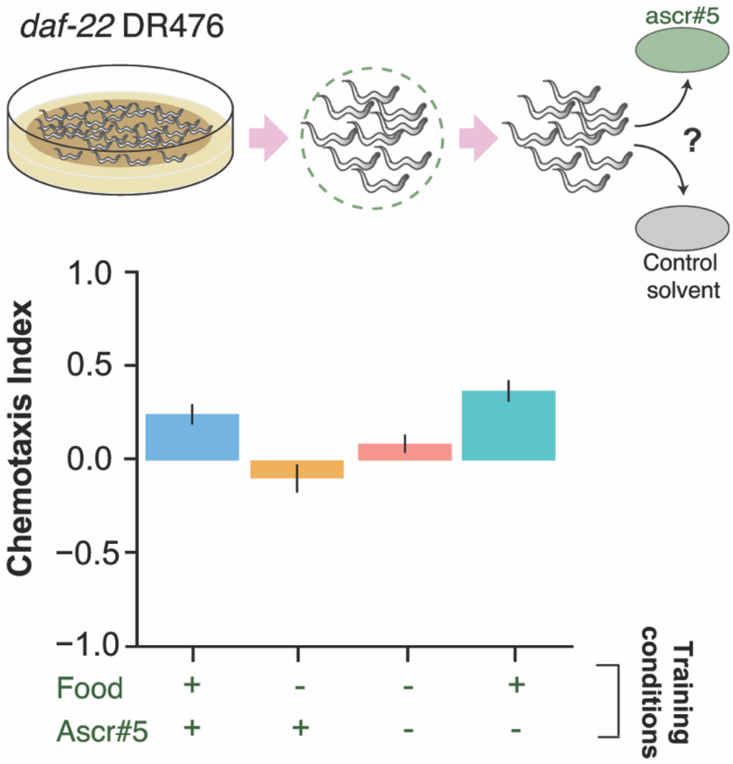


**Fig. S3.** Associative learning can reverse the response to ascr#5 of a strain that does not produce pheromones (*daf-22* DR476). Worms grow at high density and with plenty of food until young adult. Animals are then transferred to conditioning plates, where they spend 6 hours training with ascr#5. Worms are then assayed for chemotaxis to the ascaroside they were trained with. Chemotaxis index (mean CI ± SEM, n. experiments =1) for the different training conditions (indicated in green along the x axis, all at high worm density): '+ food + ascr#5' (blue bar, positive association: food and ascaroside), '–food + ascr#5' (yellow bar, negative association: starvation and ascaroside), '– food – ascr#5' (red bar), '+ food – ascr#5' (turquoise bar).

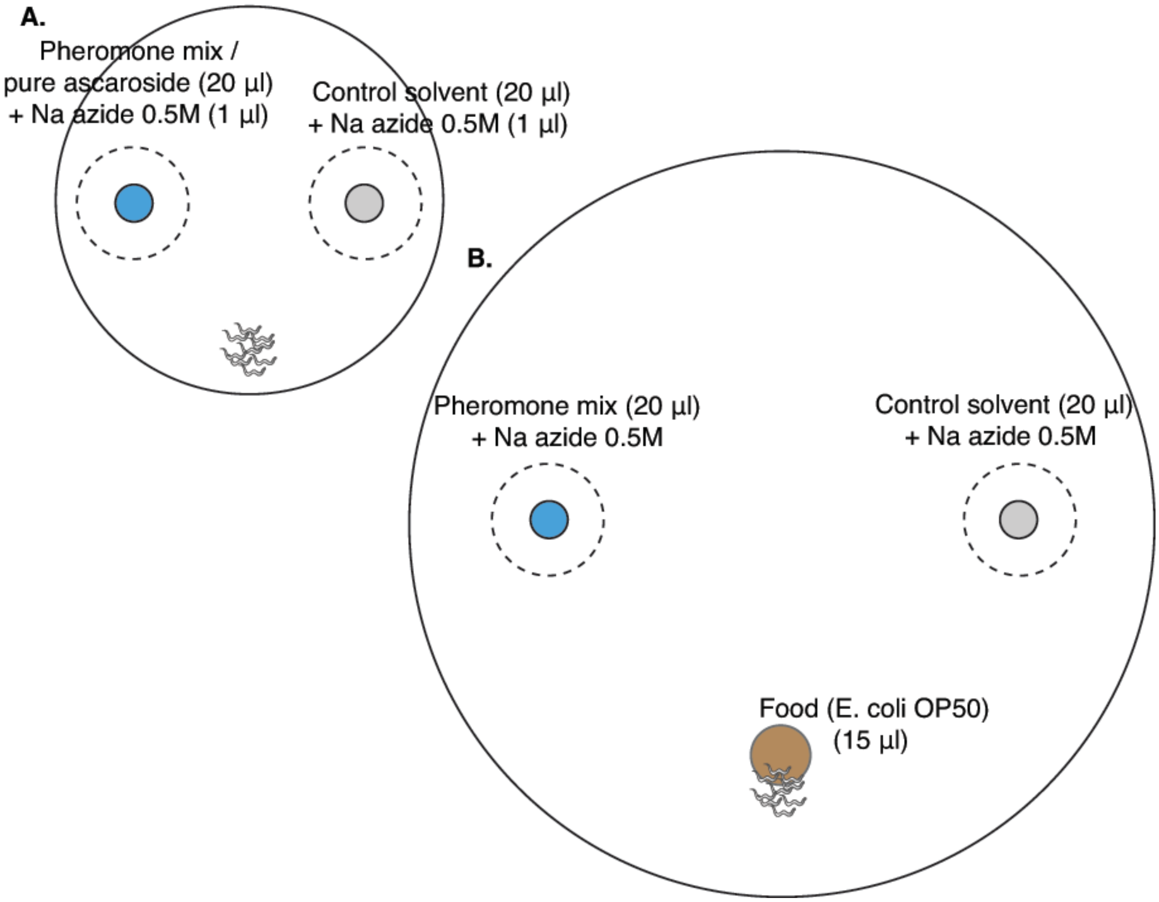


**Fig. S4.** Layout of plates for chemotaxis assays. **A**. Chemotaxis assay: 60 mm petri dish filled with 10 ml of NGM. In blue, the spot with 20 µl of either pheromone blend or pure synthetic ascarosides; in grey the spot with 20 µl of control solvent. In both spots, 1 µl of Na azide 0.5M is added to paralyze the worms once they reached the cue. Worms are placed equidistant from the two spots. **B**. Modified chemotaxis assay (choice after food): 100 mm petri dish filled with 25 ml of NGM. In blue, the spot with 20 µl of the pheromone blend, in grey the spot with 20 µl of the control solvent and in brown the food patch, 15 µl of a diluted OP50 *E. coli* culture. In the pheromone and control spots, 2 µl of 0.5 M sodium azide was added in order to anaesthetize the animals once they reached the spots. Since the anesthetic action of sodium azide lasts for about 2 hours in this set-up, another 1µl was added two hours after the beginning of the assay in both spots.
